## Supplementary figures and images for "Mechanism of outer membrane destabilization by global reduction of protein content"

### Supplemental Figure 1

#### Supplemental Figure 1

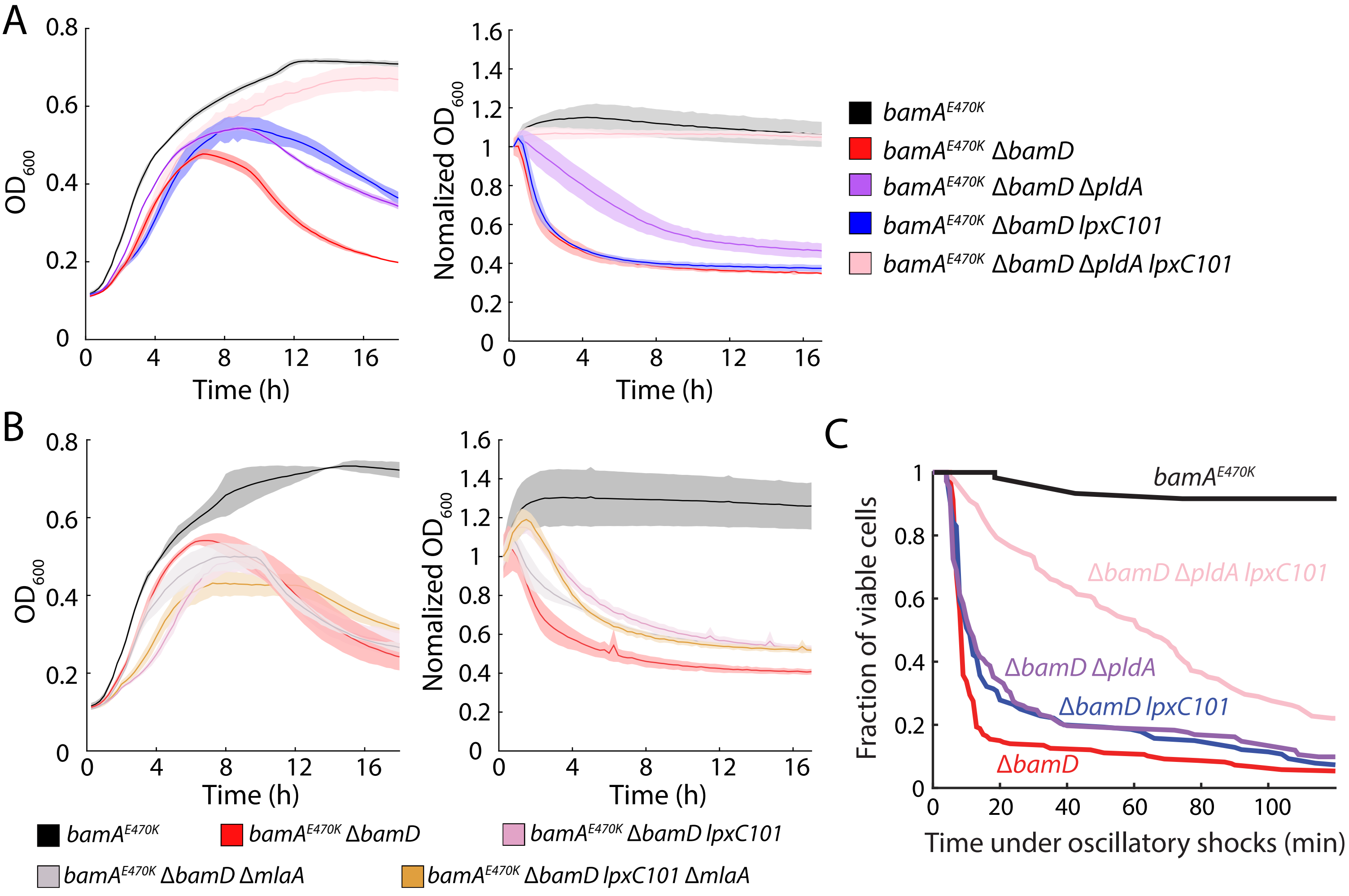
